# Click-qPCR: an ultra-simple tool for interactive qPCR data analysis

**DOI:** 10.1101/2025.05.29.656779

**Authors:** Azusa Kubota, Atsushi Tajima

**Affiliations:** Department of Bioinformatics and Genomics, Graduate School of Advanced Preventive Medical Sciences, Kanazawa University, Kanazawa, Japan

**Keywords:** Quantitative real-time PCR (qPCR), Gene expression analysis, Copy number variation analysis, Bioinformatics, Shiny web application

## Abstract

Real-time quantitative PCR (qPCR) is a pivotal technique for analyzing gene expression and DNA copy number variation. However, the limited availability of easy-to-use software tools remains a significant challenge for experimental biologists without programming expertise. To address this issue, we developed Click-qPCR, a user-friendly, web-based Shiny application that requires no local installation. Click-qPCR streamlines ΔCq and ΔΔCq calculations using user-uploaded CSV data files. The application allows users to dynamically select genes and groups, and to perform Welch’s *t*-tests for pairwise comparisons and one-way ANOVA with Dunnett’s post-hoc test for multi-group comparisons. Results are immediately visualized as interactive bar plots (mean ± SD with individual data points) and can be downloaded as publication-quality images, along with statistical summaries. Click-qPCR enables researchers to efficiently process, interpret, and visualize their qPCR data regardless of programming experience, thereby accelerating data processing and improving the accessibility of routine analysis. The Shiny application of Click-qPCR is freely available at https://kubo-azu.shinyapps.io/Click-qPCR/. The source code and user guide for Click-qPCR are openly available on GitHub at https://github.com/kubo-azu/Click-qPCR.

## Background

Real-time quantitative PCR (qPCR) is a cornerstone experiment in molecular biology, offering a highly sensitive and widely adopted method for profiling gene expression levels [1]. Its application currently spans diverse research areas, ranging from basic biological investigations to clinical diagnostics. In addition to assessing mRNA expression, relative qPCR is a robust method for detecting DNA copy number variations (CNVs), making it a common and effective technique. Although qPCR encompasses both absolute and relative quantification strategies, relative quantification is more commonly used in practice. This preference largely stems from the inherent complexities and specific requirements of absolute quantification, which often limit its utility to specialized applications, such as virus quantification. The ΔΔCq method, which is common for relative quantification, robustly compares mRNA transcript levels and CNVs. This comparison relies on a two-step normalization: first against a stable reference gene, and then against a control or calibrator sample [2]. This normalization process is crucial for accurately interpreting biological changes in gene expression and CNV analysis.

Despite its widespread adoption and critical importance for relative gene expression and CNV analysis, the ΔΔCq method often presents significant hurdles for researchers, particularly those in wet-lab environments, in terms of accessing and utilizing appropriate analysis software. Many existing software solutions are proprietary and commercially licensed, platform-specific, or demand a level of computational proficiency and programming knowledge that can be a barrier for many experimental biologists. For instance, spreadsheet-based qPCR analysis is often error-prone and time-consuming for multiple genes, while advanced statistical software can be unnecessarily complex for routine tasks.

Recognizing these limitations, we developed Click-qPCR. Unlike many existing solutions, which may require local software installation, command-line operations, or a foundational understanding of programming environments, Click-qPCR is a fully web-based application that requires no setup beyond a standard internet browser. This tool is designed to provide a user-friendly, intuitive, and readily accessible web-based platform specifically designed for the rapid and interactive analysis of qPCR data. Its focus on operational simplicity and immediate visual feedback makes Click-qPCR ideal for researchers, students, and technicians with limited bioinformatics experience.

## Implementation

The Click-qPCR application was developed using the R programming language [3] and the Shiny framework [4] to provide an interactive web interface. It relies on several publicly available R packages from the CRAN repository for its functionality. The application is platform-independent and can be run on any operating system with a standard R installation. Alternatively, users are required to have the following packages installed, which can be obtained using install.packages():

– shiny: Provides the core framework for the interactive web application [4]
– dplyr: Utilized for data manipulation, filtering, and summarization operations [5]
– tidyr: Assists in reshaping and tidying datasets for analysis [6]
– ggplot2: Handles graphical rendering of bar plots and error bars [7]
– DT: Integrates dynamic, interactive HTML tables for summary output display [8]
– RColorBrewer: Used for selecting and applying color palettes to the plots [9]
– fontawesome: Enables the use of ‘Font Awesome’ icons within the user interface [10]
– multcomp: For performing the Dunnett’s post-hoc test [11]

The application’s modular design supports both ΔCq-based relative quantification and ΔΔCq analysis workflows. All components are reactive and update automatically in response to user input, enhancing the fluidity and intuitiveness of the analytical experience. The graphical overview of Click-qPCR is in Figure 1.

**Figure 1.**
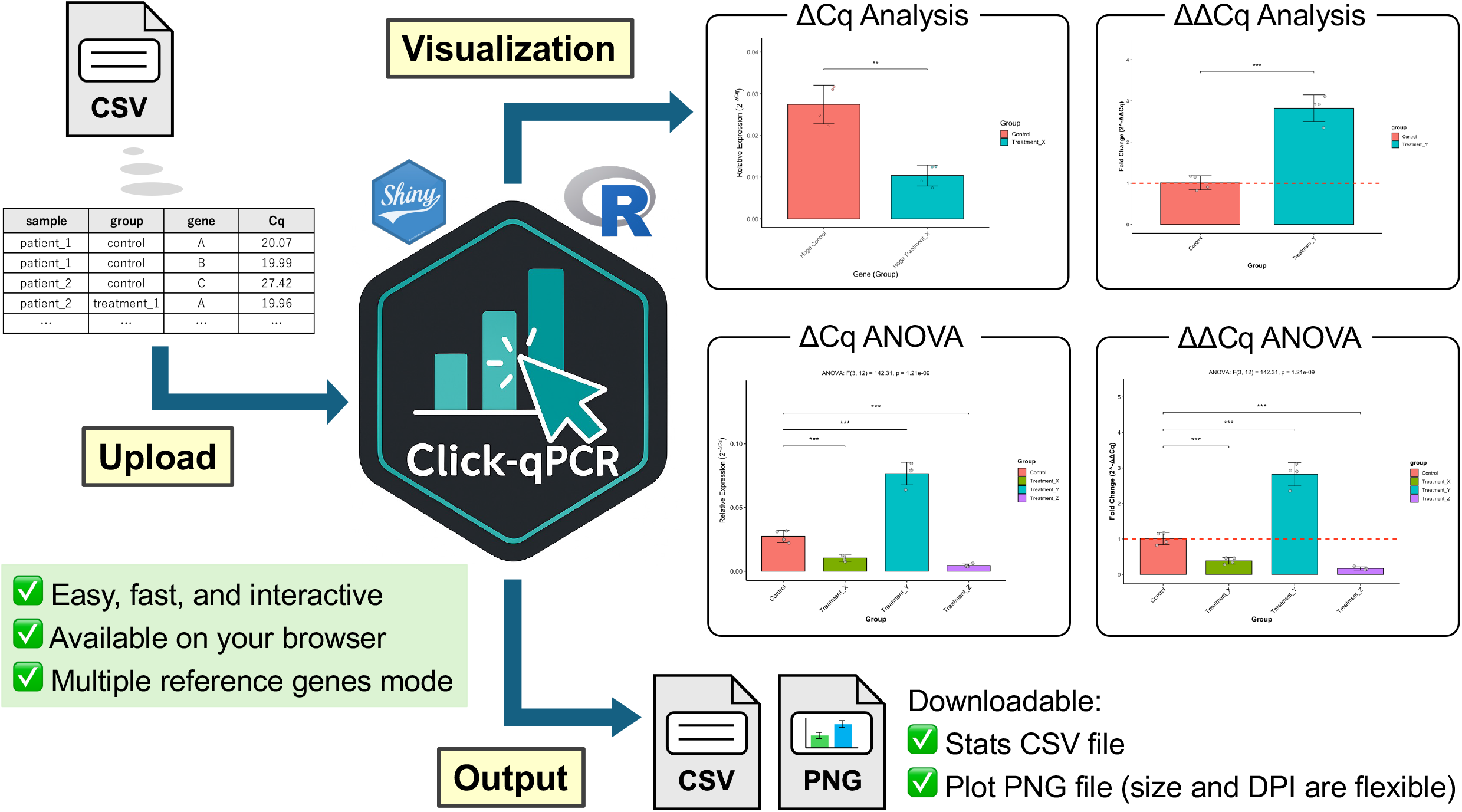
Graphical overview of the Click-qPCR application.

## Input and user interface

Click-qPCR employs a streamlined, tab-based interface comprising “Preprocessing and ΔCq Analysis”, “ΔΔCq Analysis”, “ΔCq ANOVA (Dunnett’s post-hoc)”, “ΔΔCq ANOVA (Dunnett’s post-hoc)”, and “Diagnostics” sections. The analysis process begins on the “Preprocessing and ΔCq Analysis” tab, where a tidy CSV file containing four essential columns of sample, group, gene, and Cq is uploaded. A template file and an example dataset are available for download to facilitate proper formatting. Data loading is implemented as a deliberate two-step process. Upon file selection, the first 10 rows are displayed in a sidebar preview. The file is formally loaded and validated by clicking the “Load File” button.

Within the “Preprocessing and ΔCq Analysis” tab, one or more comparisons can be defined by checking the “Enable multiple reference genes” box. This mode is available to follow the MIQE 2.0 guidelines about the number of reference genes [12]. Reference and target genes are selected via the drop-down menus, and experimental group pairs are specified for comparison. Additional comparison sets can be added via the “Add” button and deleted via the “Remove” button. The “Reset All” button restores the application to its initial state. In the “ΔΔCq Analysis” tab, the reference gene from the main analysis is retained automatically. A single target gene, a base (control) group, and one or more treatment groups are selected to compute fold changes relative to the base group.

For multi-group comparisons, users can navigate to the “ΔCq ANOVA (Dunnett’s post-hoc)” tab. Here, they select a single target gene, a control group, and two or more treatment groups. After running the analysis, the results are presented as relative expression (2^−ΔCq^). The same statistical results can also be visualized as fold-change (2^−ΔΔCq^) on the dedicated “ΔΔCq ANOVA (Dunnett’s post-hoc)” tab.

## Data processing and statistical analysis

For the ΔCq analysis, users can select one or multiple reference genes. The ΔCq value for each sample is calculated using the Cq of the target gene against the mean Cq of the selected reference gene(s). Relative expression is calculated as:

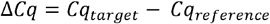

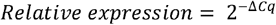

Welch’s *t*-tests are performed on the log2-transformed ΔCq values for each specified group pair. Cases with zero variance are identified, and “Zero variance” is reported in place of a test statistic. In the ΔΔCq analysis, ΔCq values are first obtained as above. The ΔΔCq and fold change values are computed as:

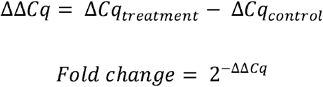

Statistical significance between treatment and control group is determined using Welch’s *t*-tests on ΔCq values.

For comparing three or more groups, the application performs a one-way analysis of variance (ANOVA) followed by Dunnett’s post-hoc test to compare each treatment group against a single control group. This analysis is performed on the relative expression (2^−ΔCq^) values.

## Visualization and output

The application adjusts plot width to accommodate the number of bars, introducing a horizontal scrollbar when necessary. Plot height is similarly expanded based on the number of significance brackets, ensuring adequate spacing between them. All analysis modes generate high-quality bar plots, displaying mean ± standard deviation (SD) with individual data points overlaid via jitter plots. Statistical comparisons are annotated with significance brackets. Significance levels are denoted as *p* < 0.05 (*), *p* < 0.01 (**), *p* < 0.001 (***), and *ns* otherwise. A corresponding summary table provides mean, SD, sample size (N), *p*-value, and the assigned significance symbol. Download options include three buttons:

– The “Download Plot” button outputs a PNG file with custom width, height, and DPI specified via sliders.
– The “Save Displayed Size” button outputs an on-screen plot PNG file.
– The “Download Stats” button generates a CSV file containing the on-screen summary table data.

## Reproducibility and diagnostics

An embedded example dataset and a downloadable template CSV file are provided to support reproducibility. The “Diagnostics” tab allows verification for core functions in any execution environment. Time-stamped downloads facilitate record-keeping. Upon clicking the “Run Diagnostics” button, the application uses the built-in dataset (similar to the template CSV file contents) to test three functions:

– First, the sample data loading step confirms successful file loading.
– Second, the ΔCq analysis validation step confirms that ΔCq calculation and *t*-tests detect the expected difference.
– Third, the ΔΔCq analysis validation step confirms that ΔΔCq (fold change) calculation and
– *t*-tests detect the expected difference.
– Fourth, the ANOVA and Dunnett’s test validation step confirms that the multi-group comparison functionality is operating correctly.

If all tests return “Passed” status, the application’s calculation and statistical processing capabilities are considered verified.

## Use case

Click-qPCR is designed to streamline relative quantification analysis for researchers working with qPCR data, particularly those without a strong background in programming or statistics. The tool is intended for diverse user groups including:

– Experimental biologists conducting small-to medium-scale gene expression studies
– Technical staff responsible for routine data analysis in core facilities
– Undergraduate and graduate students learning about qPCR and statistical evaluation
– Collaborative research groups where ease of data sharing and reproducibility are essential

## Limitations

While Click-qPCR provides an accessible and efficient platform for relative quantification analysis, several limitations should be noted:

– About assumption of normality: The app performs Welch’s *t*-tests without explicitly confirming equal variances.
– Normalization with multiple reference genes: The application now supports normalization using the geometric mean of multiple reference genes, enhancing the robustness of the analysis. However, automated methods for identifying the most stable reference genes, such as geNorm [13], Normfinder [14], and BestKeeper [15], can be conducted outside this tool.
– Simplified statistical options: The app does not offer a correction for multiple comparisons (e.g., Bonferroni correction or FDR), which may be relevant when analyzing many target genes simultaneously.
– No support for technical replicates handling: The input format assumes that users have averaged technical replicates. Raw Cq values across technical replicates must be prepared in advance.

## Concluding remarks

Click-qPCR is a web-based application designed to facilitate the rapid and reproducible analysis of qPCR data. In contrast to conventional tools that require programming skills or complex local installations, it is accessible directly through a web browser, enabling use by researchers, students, and technical staff with limited bioinformatics experience. The application supports multiple primary workflows of relative expression analysis: ΔCq analysis for direct group comparisons using Welch’s *t*-tests, ANOVA with Dunnett’s post-hoc test for multi-group comparisons, and ΔΔCq analysis for calculating fold changes relative to a control group. It generates publication-ready visualizations and includes a dedicated Diagnostics tab for verification of core computational functions. By integrating a guided interface, flexible export options, and reproducibility-oriented features, Click-qPCR enables efficient processing of qPCR datasets and promotes transparent reporting of gene expression analyses.

## Supporting information

Template CSV file for Click-qPCR

## Declarations

### Availability of source code and requirements

– Project name: Click-qPCR
– Project home page: https://kubo-azu.shinyapps.io/Click-qPCR/ (Shiny); https://github.com/kubo-azu/Click-qPCR (R source code)
– Operating System(s): Platform Independent
– Programming language: R
– License: MIT

### Data availability

The template CSV file for Click-qPCR can be found in the Supplementary information and at https://github.com/kubo-azu/Click-qPCR/blob/main/Click-qPCR_template.csv.

## Acknowledgments

Writing - original draft: AK. Writing - review and Editing: AT. Conceptualization: AK and AT. Software: AK. Funding Acquisition: AK. Supervision: AT.

This work was supported by JST SPRING, Grant Number JPMJSP2135. The authors thank Dr. Yasuhito Shimada and Dr. Liqing Zang for their constructive feedback, which contributed to the improvement of Click-qPCR.

## Ethical considerations

The authors declare that ethical approval was not required for this type of research.

